# Multi-omics guided pathway and network analysis of clinical metabolomics and proteomics data

**DOI:** 10.1101/2025.06.26.661095

**Authors:** Christina Schmidt, Thomas Naake

## Abstract

Metabolomics, the study of small molecules in biological systems, is a powerful tool for understanding biochemical pathways, discovering biomarkers, and elucidating disease mechanisms. This chapter provides a guide to performing metabolomics data analysis in *R*, focusing on enrichment analysis and network-based approaches. It covers essential steps in data processing, quality control (QC), differential expression analysis, integration with proteomics using multi-omics factor analysis (MOFA) and statistical network analysis, as well as enrichment analysis using prior knowledge. The methods outlined provide a framework for biomarker discovery and advancing systems-level understanding of disease processes using metabolomics data in combination with prior knowledge and proteomics data.

## Introduction

With the increasing number of metabolomics studies, standardised and reproducible analysis strategies are essential to ensure accurate biological interpretation. To support this, high-quality prior knowledge (PK) resources, such as curated metabolite sets and gene-metabolite associations, combined with the application of metabolic networks are effective to facilitate data interpretation. Networks provide a powerful framework to study metabolomics and enable the integration with proteomics data by representing metabolites and protein as vertices and their relationships (e.g., correlations, biosynthetic enzymes) as edges.

This approach enables insights into system-level changes and interactions between metabolites, proteomics and other omics layers.

In this chapter, we will showcase the statistical programming language R’s extensive ecosystem for metabolomics analysis and integration with proteomics, PK using network analysis among other strategies, including packages like *igraph* (Csardi and Nepusz 2006), *MOFA* (Argelaguet et al. 2018), *decoupleR* (Badia-i-Mompel et al. 2022) and *MetaProViz* (Schmidt et al. 2025). As an example of multiple data modalitie integration, we integrate metabolomics with proteomics that share the same sample set using Multi-omics factor analysis (MOFA), a statistical technique to identify latent factors of shared variation in multi-omics datasets. We use the results from the MOFA model to guide downstream analysis in pathway and network analysis. *MetaProViz* provides access and quality control plots for the integration of metabolomics with prior knowledge, whilst *decoupleR* uses those biological signatures to calculate scores based on prior knowledge, offering the interpretation of the datasets by known biological networks and pathways.

In this chapter, we distinguish between knowledge and experimental networks (Amara et al. 2022). Knowledge networks are constructed using existing biochemical and biological information, helping to interpret metabolomics and proteomics data within the framework of known pathways. For example, a metabolic reaction network represents a knowledge network, where metabolites act as vertices and their biochemical conversions are the edges. In contrast, experimental networks are directly derived from metabolomics and proteomics data, linking metabolites and proteins based on observed relationships like correlation or, in case of MS/MS metabolomics data, spectral similarity. Both types of networks can be explored using advanced statistical methods, graph analysis, and data-driven techniques to reveal meaningful patterns and connections within the dataset.

## Materials

We will use the dataset from Gegner et al. (2024) to showcase functionality in *R* to analyse metabolomics and proteomics using network and pathway analysis tools. We present here an end-to-end pipeline starting with data import, data processing, quality assessment and control, multi-omics integration and downstream analysis to interpret the latent factors of the model. The datasets are proteomics and metabolomics measurements from fresh-frozen tumor and non-tumorous (adjacent) tissue of patients with lung adenocarcinoma. Metabolomics measurements were acquired using the MxP® Quant 500 kit (Biocrates) using LC-MS/MS and FIA-MS/MS measurements run on a UPLC I-class PLUS (Waters) system coupled to a SCIEX QTRAP® 6500 + mass spectrometry system in electrospray ionization (ESI) mode. Proteomics measurements were aquired using an Easy-nLC^TM^ 1200 system (Thermo) coupled to a timsTOF Pro mass spectrometer (Bruker Daltonics). For further details on the analytical methods refer to Gegner et al. (2024).

Due to space constraints, the code for all analyses will not be displayed in this chapter. Interested readers are refered to the complete and reproducible protocol available at www.github.com/tnaake/MiMB_networks/R/chapter_network_multiomics.qmd.

## Methods

### Data processing and quality assessment and quality control (QA/QC)

Processing and quality assessment and quality control (QA/QC) are critical to ensure the integrity of metabolomics data. Steps typically include detection control of outliers, removal of batch effects, data normalization and transformation, and missing value imputation (optional). Each step requires careful parameter selection based on the dataset and experimental conditions. In the case of multi-omics analysis using MOFA, as presented here, it is paramount to ensure that all modalities share the same set of samples, with identical sample names and order, to facilitate accurate integration and analysis.

#### Load the datasets, data wrangling

1. Load the proteomics and metabolomics datasets via import functions from *MatrixQCvisUtils* (available via www.github.com/tnaake/MatrixQCvisUtils). Importing the XLSX sheets into *R* via the *MatrixQCvisUtils* package, creates *SummarizedExperiment* (Morgan et al. (2024)) objects. The *SummarizedExperiment* class stores the measured data (available via *assay*), next to the metadata associated to the samples (available via *colData*) and features (available via *rowData*).
2. After importing the datasets, check the following properties:

- correct representation of missing values (*NA*) in dataset,
- correct data dimension and structure,
- consistent number of samples and matching sample names between datasets,
- verify that the sample order is identical across the modalities,
- correct formatting of feature names, complete number of features for each dataset,
- complete and correctly formatted metadata associated to each dataset
3. (optional) Depending on 2., identify the nature of missing features, harmonize sample names, correct feature names, harmonize metadata and check for completeness.
4. (optional) Identify additional metadata, for example clinical parameters or translated IDs for metabolites (e.g. HMDB, KEGG, ChEBI ids) and proteins (e.g. Entrez, SYMBOL, Uniprot ids), from other sources. Add the metadata to the respective slots of the *SummarizedExperiment* object: *rowData* for feature-related and *colData* for sample-related metadata).

#### Filtering of features in the datasets

Each dataset includes typically information on the reliability of features, which can be used to remove non-reliable features. This step improves the quality of the datasets. Filtering and removal of the features should be tailored to the study design and the characteristics of the dataset. While stricter filtering can reduce false positives, it may also remove true low-abundance features.

1. Remove low or high abundance features. Low intensity features may be due to noise or below the limit of detection or quantification. High intensity may be above the limit of quantification, the linear dynamic range, or the calibration range. Removing these features prevents artifacts in downstream statistical analyses. For the targeted metabolomics dataset, ensure that the dataset only includes high-quality metabolite measurements. The MetIDQ output provides assessment of the quality of each measurement, encoded by a color code of each cell (dark blue: < LOD (Limit of Detection; 3x signal to noise); light blue: < LLOQ (Lower Limit of Quantification; 10x signal to noise) or > ULOQ (Upper Limit of Quantification); green: valid; Yellow: Internal Standard out of range). Remove the low-quality features that do not meet the defined quantification criteria (*choice of user*). In this case, we will retain only those metabolites where at least 50% of the measurements fall within the quantification limits (valid). This filtering step improves data reliability and minimizing the impact of imprecise data points. Other filtering steps can also be applied, e.g. the 80% filtering rule or adapted-80% filtering rule Wei et al. (2018)
2. Verify that all measurements fall within expected ranges (e.g., nonnegative values for intensities). For measurements that fall outside the expected range, correct them, e.g., by setting these values to *NA* (missing data).
3. (optional) Remove those features that are only detected in a subset of samples (*choice of user*). High levels of missing data (*NA*) values can reduce statistical power. While imputation techniques can be applied, imputation of excessive numbers of missing values per feature can lead to unreliable results. Additionally, features identified in only a few biological replicates may lack reproducibility, impacting downstream analyses. Filtering criteria can be set on a global level, such as removing features absent in more than 50% of the samples, or based on metadata, such as removing features detected in fewer than two biological replicates per condition. More stringent thresholds, such as requiring detection in at least 50% of replicates within each condition, can further improve dataset reliability. The choice of filtering thresholds should be guided by domain-specific knowledge and the characteristics of the datasets. Overly strict filtering may remove biologically relevant proteins that are present only in specific subsets of samples, such as particular cell lines or patient groups. Filtering strategies should be carefully considered to maintain a balance between data quality and the retention of meaningful biological information.
4. Remove the features with high coefficient of variation (CV, *choice of user*). High variability in technical replicates or QC pools suggests inaccurate quantification. For example, features with CV > 30-50% of raw intensities across technical replicates may be removed.
5. (optional) Remove features with a standard deviation of 0 as these features do not vary across samples and do not contain information for biological interpretation.
6. For proteomics data, ensure that the dataset only includes high-quality protein measurements. In the proteomics dataset generated by MaxQuant (Cox and Mann 2008), peptide counts serve as an indicator of the reliability of protein identification. To ensure data quality, we will filter out proteins identified with low confidence (*choice of user*). In this case, we will remove any protein that has fewer than two peptides supporting its identification. This approach helps retain only well-supported protein identifications, enhancing the robustness of downstream analyses.
7. For proteomics data, remove common contaminants by using a contaminant database (e.g., MaxQuant’s contaminant list). Contaminants like keratins (from human skin), trypsin (from digestion), or bovine serum albumin (BSA) can interfere with interpretation of results.
8. For the proteomics dataset, remove the features with reverse sequences as they do not correspond to real biological proteins. Reverse sequences refer to decoy sequences generated by reversing the amino acid sequence of real proteins. Reverse sequences are commonly used as negative controls in target-decoy strategies for false discovery rate estimation in mass-spectrometry-based proteomics.

#### Quality assessment and quality control (QA/QC)

The next step in the processing pipeline is to assess the quality of the data, including both the measurements and the metadata values, and to control for it.

The *MatrixQCvis* package (Naake and Huber 2022) performs QA/QC on *SummarizedExperiment* objects and offers users to perform these steps in an interactive *shiny* application.

1. Launch the application via *shinyQC(metabolomics)* or *shinyQC(proteomics)* to begin the quality assessmet for the respective datasets.
2. Check the number of measured and missing values per sample and per feature. The metric informs about the dynamic range of the acquisition. Differences between samples of an experiment may indicate differences in the dynamic range and/or in the sample content.
3. Check the distribution of measurements per sample. The metric informs about the dynamic range of the acquisition. Differences between samples of an experiment may indicate differences in the dynamic range and/or in the sample content.
4. Examine the mean-sd plot (Figure 1 A and B). The metric informs about the level of homoskedasticity of the dataset which may be important for parametric tests. Homoscedasticity is an assumption of parametric tests, e.g. t-tests. In case of sd-mean independence, the running median should be approximately horizontal.
5. Examine the MA plots and Hoeffding’s D statistic plot. The metric informs about systematic biases and variability in the data by taking into account the log2-fold change between two conditions (*M = log2(I_i) - log2(I_j)*) and the mean expression of two conditions (*A = 1/2 (log2(I_i) + log2(I_j))*), where *I_i* can be the intensity from one sample i and *I_j* the averaged intensities from a set of samples excluding sample *i*. Hoeffding’s *D* statistic measures the dependency between *A* and *M*. It is a measure of the distance between *F(A, M)* and *G(A)H(M)*, where *F(A, M)* is the joint cumulative distribution function (CDF) of *A* and *M*, and *G* and *H* are marginal CDFs. The higher the value of *D*, the more dependent are *A* and *M*.
6. Examine the empirical cumulative distribution function (ECDF) plots. The metric informs about the distribution, the skewness of a sample and helps to identify outliers.
7. Examine the distance matrix between samples (Figure 1 C and D). The metric informs how similar or different samples are from one another based on the measured values. It informs about relationships and consistency between samples of the same group (e.g. treatment vs. control). The metric also helps to detect outliers and to identify batch effects.
8. Examine dimension reduction, e.g. principal component analysis (PCA, Figure 1 E and F) and loadings plot. PCA preserves the variance between data points and aids in exploring relationships between samples and features. PCA is helpful in identifying structure in the dataset and in detecting outliers. Make sure that pool samples cluster at the zero-zero axes of a PC1-PC2 plot.
9. Identify and remove any outliers based on the steps above. Reassess the data after the removal by repeating the steps 1.-8.
10. After reviewing the metrics 3.-9., identify the most appropriate methods for normalization, batch correction, transformation, and imputation for your datasets to ensure consistency and quality across samples and features.

**Figure 1:**
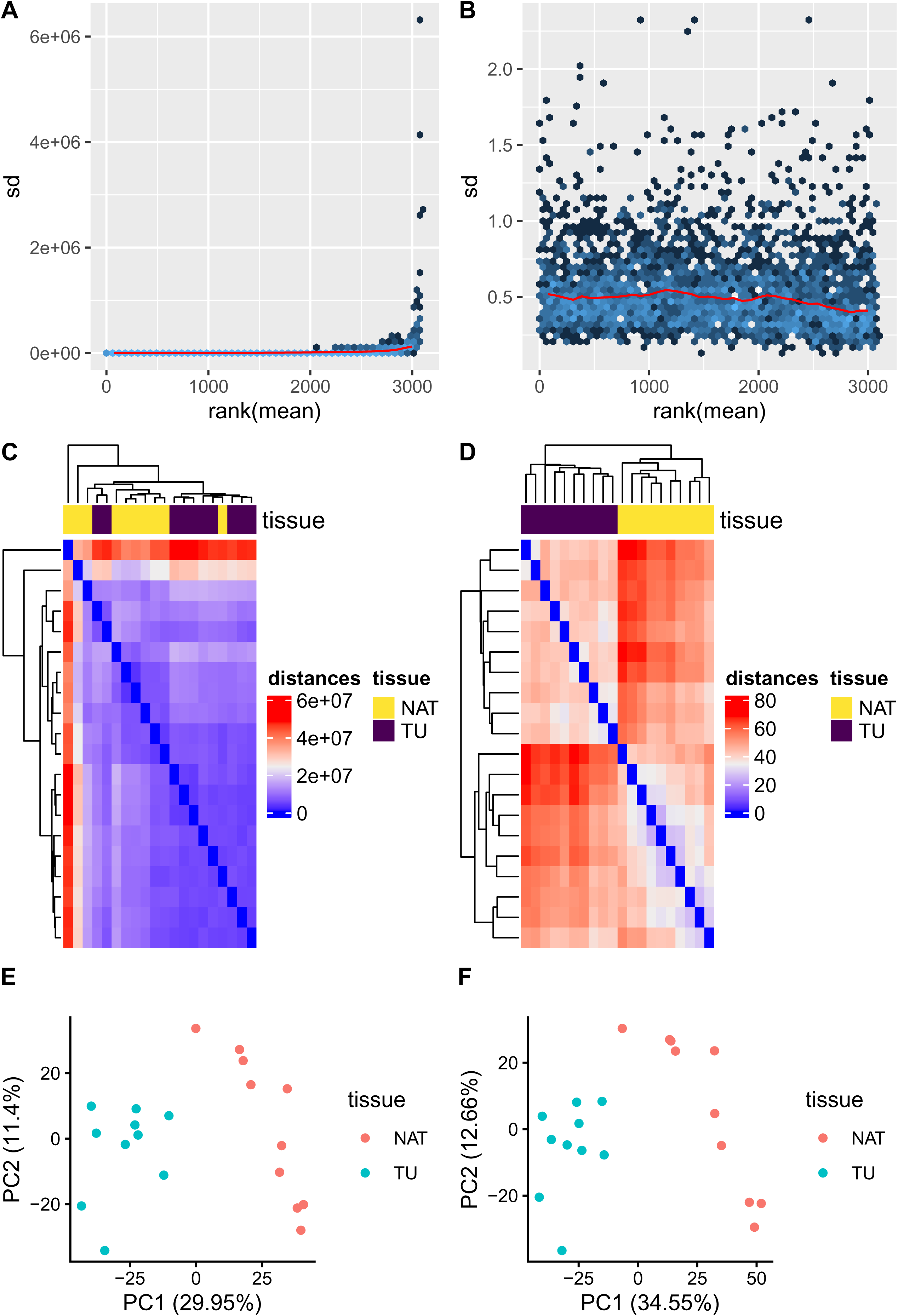
Assessment of log-transformation of proteomics dataset on quality metrics. A: Mean-sd plot of raw data. B: Mean-sd plot of transformed data. Log-transformation controls heteroskedasticity. C: Matrix of Euclidean distances between samples of raw data. D: Matrix of Euclidean distances between samples of transformed data. E: PCA of raw data. F: PCA of transformed data. Log-transformation increases variation on prinpical component 1 and 2. The percentages indicate the explained variance per principal component. PC: principal component. sd: standard deviation.

#### Normalization, batch correction, transformation, missing value imputation

Tools, such as *MatrixQCvis* (Naake and Huber 2022) can be used to explore data quality interactively and to optimize the choice of batch correction, normalization, or transformation methods. The *MatrixQCvis* package offers utility functions to perform normalization, batch reduction, transformation and imputation using a variety of methods (*normalizeAssay*, *batchCorrectionAssay*, *transformAssay*, *imputeAssay*).

1. (optional) Normalize the dataset. Normalization adjusts for technical variability and ensures comparability of metabolite and protein levels across samples. The choice of method depends on the dataset’s variability and experimental design. Common methods are median normalization, quantile division normalization (division of measurements of each sample by the given quantile, e.g. 50% for median division), sum normalization (division of measurements of each sample by the sum of intensities column for that sample).
2. (optional) Correct for batches. Batch effect correction is typically essential in multi-batch experiments acquired over a longer time, but may be not needed in smaller scale experiments. *ComBat*, for instance, requires specifying batch labels and optionally including covariates to retain biological variability. In case of linear effects of batches the batch effect can be removed via *limma*’s *removeBatchEffect* function and specifying the batch labels.
3. Transform the dataset. For example, *log* transformation is commonly used to reduce skewness and heteroskedasticity, other methods, such as *log2*, *log10*, or variance stabilizing normalization (vsn) can be applied as well. After transformation, check the mean-sd plot to assess homoskedasticity and determine if the transformation has balanced the variance.
4. (optional) Scale the features. Scaling standardizes the magnitude of intensities across features, making them comparable by putting them on a common scale (e.g., z-score).
5. (optional) Impute the missing values (*NA*) of the datasets. Metabolomics and proteomics data acquired by mass spectrometry typically contain missing values. The choice of the imputation method depends on the mechanism of missingness (e.g., missing completely at random, missing at random, missing not at random). Various imputation techniques are available in *MatrixQCvis*, such as k-nearest neighbors imputation or imputation by minimum values. For a more in-depth discussion on imputation methods in metabolomics, refer to Wei et al. (2018), and for proteomics, see Lazar et al. (2016). Generally, missing values should be imputed with caution, as doing so requires assuming a model of missingness and can introduce bias. In many cases, downstream analyses can tolerate a certain level of missing values without significant impact on results. Tools such as *limma* (Ritchie et al. 2015) and *MOFA* are designed to handle datasets with incomplete observations, making imputation unnecessary for many applications.

### Multi-omics integration using MOFA

MOFA (Argelaguet et al. 2018) integrates omics modalities using a factor analysis to uncover latent factors that explain the shared variation across different datasets. Proper data wrangling and processing as discussed in previous sections (e.g., normalization, transformation) are critical steps in ensuring the reliability and interpretability of the results.

#### Model building

1. Define the modalities (omics data types) to be included in the model. Make sure that the modalities are properly normalized and transformed. (optional) Define group information. Group information refers to the different subsets or categories within the data that can be used to model variation across multiple omics layers. This could represent different experimental conditions, sample types (e.g., tumor vs. non-tumor), or other biological categories that might drive the variation in the data.
2. Define the parameters for training of the MOFA model (*choice of user*). Parameters control how data is preprocessed before training, such as whether views or groups are scaled to the same variance or centered (with a mean of zero). Parameters can define the structure of the MOFA model, such as the number of latent factors to infer, which can be guided by prior biological knowledge or model selection techniques. The user can also specify if a spike-and-slab prior on factor loadings is applied, which promotes sparsity in factor loadings by ensuring that only a few features have strong contributions while others are near zero. The user may apply the automatic relevance determination (ARD) priors on factor loadings, allowing the model to automatically determine which factors are relevant for each view. In addition, parameters can specify the training process of the model, including the mode of convergence of the algorithm, the use of stachastic inference, the seed for reproducibility, and the maximum number of iterations before the model training is interrupted.
3. Add the model parameters to the untrained MOFA model and initiate the training of the model. The model will learn latent factors that explain the shared variation across the different omics datasets, based on the specified configurations and data characteristics.
4. Inspect the sanity of the model output. Plot data overview and verify that the dimensions for each dataset are as expected. Calculate the correlation scores between the factors. The correlation coefficients between the factors should be low, indicating that each factor captures distinct variation in the data. If factors are found to be highly correlated, it may suggest that the model is not effectively separating the variation across the datasets. In such cases, consider reducing the number of trained factors. This will not only help in improving the interpretability of the model but also enhance the biological relevance of the factors by focusing on more distinct sources of variation.
5. Examine explained variance of the factors and per modality (Figure 2 A). The analysis shows the proportion of variance explained by each latent factor across different views (modalities) and groups (if applicable). Higher variance explained per factor means that the factor captures a significant portion of the variation in that particular view. If the first few factors explain most of the variance, it suggests that only a subset of factors drive the structure in the data. If certain factors explain more variance in one view compared to another, it suggests that the factor is more specific to that modality. Co-variance in most factors indicates a strong relationship between the data types. If plotted separately, differences in variance explained between groups may indicate distinct biological patterns or variation in data structure across conditions.

#### Model interpretation: Variance decomposition and analysis of factors

1. Obtain sample covariates and cofactors and add to metadata. Metadata can be retrieved from the *colData* slot of the *SummarizedExperiment* objects.
2. Analyse the association between latent factors and metadata (e.g., clinical metadata, information on demographics, experimental condition, Figure 2 B). Compute correlations between the learned MOFA factors and the user-specific covariates (continous or categorical). Visualize and examine the correlation coefficients and p-values. The intensity and sign of the color indicate the strength and direction of the correlation. Positive correlations suggest that an increase in the covariate is associated with an increase in the factor values. Negative correlations suggest an inverse relationship.
3. Plot a scatterplot of the samples by factors (Figure 2 C). Each factor arranges the samples along a one-dimensional axis centered at zero, where the absolute value is not important; only the relative positioning of the samples matters. Samples with different signs on the axis indicate opposite “effects” along the inferred variation axis, with larger absolute values suggesting stronger effects. The interpretation of these factors is similar to that of principal components in PCA, where each factor represents a direction of variation in the data, and the separation of samples along these factors provides insight into the underlying patterns.
4. Analyse the weights of the factors. The weights represent how strongly each feature is associated with each factor. Features with little or no association to a factor will have values near zero, while those with a strong association will exhibit large absolute values. The sign of the weight indicates the direction of the effect: a positive weight means the feature has higher lievels in samples with positive factor values, while a negative weight indicates higher levels in samples with negative factor values.
5. Analyse the feature intensities across samples and relationships between features using the information from the MOFA model (Figure 2 D). The visualizations are helpful for identifying patterns clusters, assessing data quality and batch effects, detecting potential outliers, and understanding how different features, factors, or sample groups contribute to the overall variation in the dataset. The visualizations should be created for each factor of interest and for the most important (e.g. top 10 or 20) features per factor (*choice of user*).

### Differential Expression Analysis

Differential analysis identifies metabolites with significant abundance changes between conditions (Huang and Wang 2022). Here, selecting appropriate statistical tests depends on the data distribution and experimental design.

**Figure 2:**
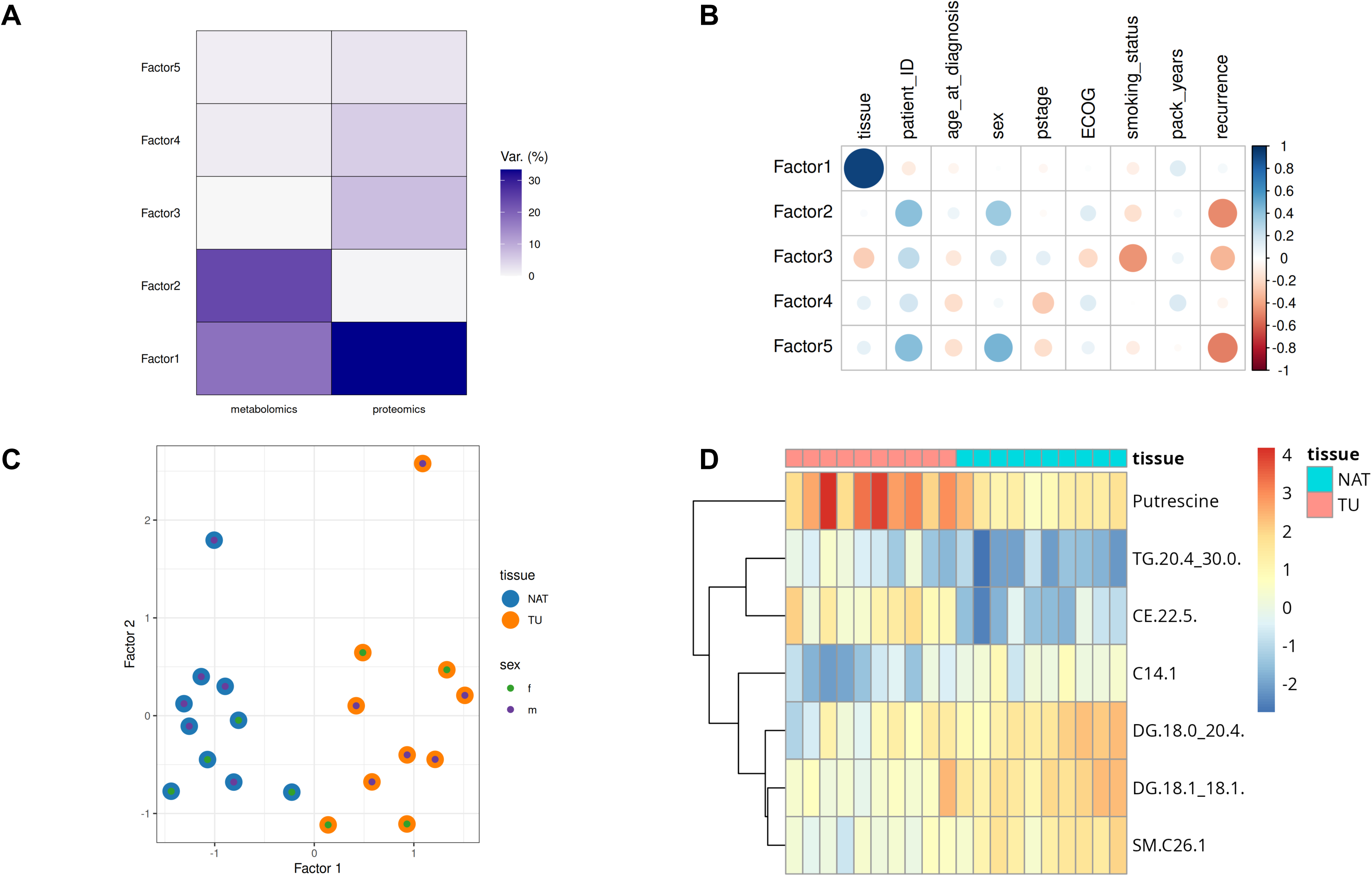
Multi-omics factor analysis diagnostic and loading plots. A: Variance explained by Factors 1-5 in the metabolomics and proteomics layers. B: Sample-based Pearson correlation coefficients between MOFA factors and meta-data variables. C: Scatterplot of Factor values for each sample for Factor 1 and Factor 2. Factor 1 separates the samples along the NAT-TU axis. D: Heatmap of transformed metabolite values for Factor 1. Shown are the top seven metabolites with the highest absolute loadings.

#### Statistical Testing

To systematically define which statistical tests to consider one can follow these steps:

1. To define which comparison(s) should be performed, it is important to understand the meta information in the data and define the biological question. For ***one_vs_one (single comparison)*** analysis, where a single condition is compared against another (e.g. numerator = “Condition1” and denominator = “Condition2”), statistical tests such as the Student’s t-test (Padhee and Ahmed 2024), Wilcoxon rank-sum test (Wilcoxon 1945), or linear regression modeling (Ritchie et al. 2015) can be used. For ***all_vs_one (multiple comparison against a reference)***, where multiple conditions are compared individually against a single reference condition (e.g., numerator = [“Condition2”, “Condition3”, …, “ConditionN”], denominator = “Condition1”) or ***all_vs_all (multiple comparison)***, where all conditions are compared against each other (e.g., numerator = [“Condition1”, “Condition2”, …, “ConditionN”], denominator = [“Condition1”, “Condition2”, …, “ConditionN”]), multiple testing has to be performed for which statistical tests such as ANOVA (analysis of variance), Welch’s ANOVA (Welch 1951), Kruskal-Wallis test (Padhee and Ahmed 2024), or linear modeling (e.g., using limma’s linear model fitting (Ritchie et al. 2015)) are appropriate.
2. Check the assumption of normality to understand if the data follow a normal- or not-normal distribution by for example performing a Shapiro test (Padhee and Ahmed 2024). For normally distributed data, t-tests or ANOVA can be used, whilst for not-normally distributed data, non-parametric tests like Wilcoxon rank-sum test or Kruskal-Wallis test are appropriate (Padhee and Ahmed 2024). The R package MetaProViz can be used to perform the Shapiro test for normality. and the Bartlett’s test for homoscedasticity
3. Check if the data are homoscedastic, meaning whether they have equal variance, using Levene’s test or Bartlett’s test (Viwatwongkasem, Vorapongsathorn, and Taejaroenkul 2004). If the data are not homoscedastic, consider using test like Wilcoxon rank-sum test since tests like Kruskal-Wallis assumes equal variance (Padhee and Ahmed 2024). MetaProViz can be used to perform the Bartlett’s test for homoscedasticity.
4. Perform differential expression analysis. If there are missing values (*NA*s) in the data linear modeling can be used, such as limma, which fits a linear model to the data and can handle missing values (Ritchie et al. 2015). Yet, the limma method assumes that the residuals of the linear model are approximately normally distributed (Ritchie et al. 2015). Other options are to either perform missing value imputation, which comes with its own pitfalls in metabolomics, or to assume that the missing values are true zeros, which in turn can lead to zero inflation (Huang and Wang 2022). MetaProViz can be used to perform differential analysis. It gives the choice between different statistical tests. Importantly, MetaProViz also adjusts for multiple testing using the Benjamini Hochberg, False Discovery Rate (FDR), Bonferroni or Holm method (Schmidt et al. 2025).

#### Visualization of differential analysis results

1. Check the results of the differential analysis by a volcano plot. Check the proportion of the metabolites that are significantly up- or down-regulated e.g. between the two conditions for both, the t-test and Wilcoxon rank-sum test (Figure 3 C-D). The classical volcano plot of the differential analysis results(Figure 3 B) displays the log2 fold change on the x-axis, while the y-axis represents the negative logarithm of the adjusted p-value. The volcano plot is a scatter plot that displays the relationship between the magnitude of change (log2 fold change) and the significance (adjusted p-value) of each metabolite. Points above a certain threshold are considered statistically significant and hence it is important to set thresholds for significance (e.g. adjusted p-value < 0.05) and log2 fold change(e.g. |log2FC| > 0.5).
2. (optional) Add additional information to the volcano plot. For example, with the *MetaProViz* package, add additional relevant information to the plot (Schmidt et al. 2025), e.g. the metabolite name and colour code for metabolite classes as provided by Biocrates (Figure 3 E).
3. (optional) Examine individual plots for each metabolite class for better interpretability (Figure 3 F-H).

### Analysis using biological prior knowledge

Prior Knowledge (PK) is a powerful tool in omics data analysis, allowing researchers to leverage existing biological information to gain mechanistic insights into their data.

1. Choose the PK that should be applied. The type of PK will dictate the biological question that can be answered and how the results can be interpreted.

- Collections of biological pathways or processes, i.e. sets of genes/metabolites with common characteristic (e.g. GO-term, KEGG,etc.), commonly also referred to as pathway analysis. Pathway analysis is used to display that the features of interest are statistically more enriched in specific pathways (García-Campos, Espinal-Enríquez, and Hernández-Lemus 2015). The dysregulation of a pathway is inferred from measurements of its own components (e.g. metabolite abundance).
- Collections of enzymes/molecules and their targets, for example, kinases and target phosphopeptides, which are usable for footprint analysis. Footprint analysis estimates the activities from molecular readouts of the targets (e.g. phosphosites) downstream of the enzyme (e.g. kinase) (Dugourd and Saez-Rodriguez 2019).
- Collections of enzymes/molecules, their targets and the interaction between them, which will depict a PK network (PKN), for which network-based approaches are needed to systematically integrate the data and extract coherent patterns (Dugourd et al. 2021).
2. Choose a statistical method to be applied on the PK taking into account the biological perspective and method performance (Badia-i-Mompel et al. 2022). In case of classical mapping and fooprint analysis, statistical methods such as Gene Set Enrichment Analysis (GSEA), Over-Representation Analysis (ORA), multivariate linear models (MLM), univariate linear models (ULM), can be used to obtain a score for the activity of a kinase (foot-printing) or the dysregulation of a pathway (mapping) (Badia-i-Mompel et al. 2022). The choice of statistical method will influence the results and the biological interpretation. For instance, GSEA uses a feature list ranked by their expression changes between conditions and hence finds pathways enriched at the top or bottom of the ranked gene list, whilst ORA is based on Fisher’s exact test and checks if a selection of upregulated features (e.g. log2FC >1, p.adj < 0.05) are over-represented in a specific pathway (Badia-i-Mompel et al. 2022). In a benchmark study evaluated on transcriptomic and phospho-proteomic perturbation, linear models (e.g. ULM, MLM, etc.) outperformed GSEA, whilst ORA performs similarly well (Badia-i-Mompel et al. 2022). Here it is also important to note that the chosen thresholds for e.g. up-regulated metabolites will affect the ORA results as it affects the basket size. The choice of ranking by log2FC versus ranking by t-value, which is a combination of change and statistics, will alter the GSEA results. Dependent on other available analysis, such as MOFA or PCA, the ranking of features can also be done using those outputs. Similarly, using the MOFA results created above, one can also define the top and bottom 20% of altered features, which in turn can be used to perfrom ORA.

**Figure 3:**
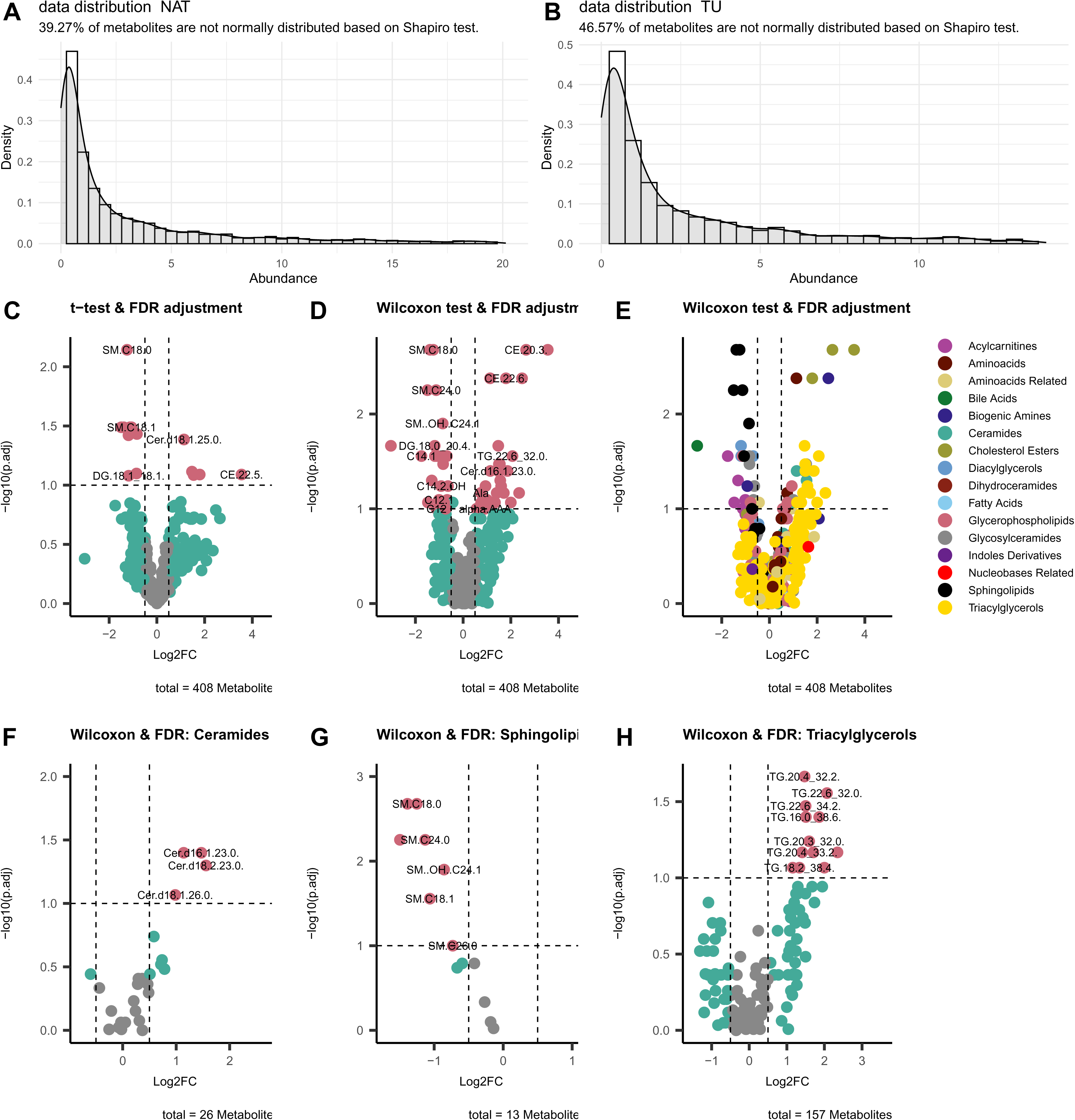
One-versus-one differential expression analysis between tumour (TU) versus normal adjacent tumour (NAT). A: Data distribution for each metabolite abundance of NAT B: Data distribution for each metabolite abundance of TU. The Shapiro test results shows that the majority of the metabolites follow mostly a normal distribution, which means it is appropriate to perform a t-test. It is important to note that almost half of the metabolites are not normally distributed, which means that the t-test may not be appropriate for these metabolites. In this case, it might be better to use a non-parametric test such as the Wilcoxon rank-sum test or Kruskal-Wallis test. C-H: Volcano plots of differential analysis comparing TU versus NAT using either the t-test and FDR adjustment (C) or the Wilcoxon rank-sum test and FDR adjustment (D-H). E is colour-coded based on the metabolite class as defined by Biocrates, whilst F-H) are selected metabolite panels based on Biocrates class.

Yet, it is relevant to decide if performing enrichment analysis will be able to answer the biological question at hand. Metabolites are not genes, which means if a pathway is up-regulated in the context of genes, all the genes involved in that pathway produce more copies, whilst metabolite levels show lower coherence during regulation since the up-regulation may increase the abundance of the end-product but not of the metabolite intermediates(Lee, Su, and Huan 2025). Hence, pathway analysis in metabolomics can produce misleading or nonsensical results due to the unique characteristics and constraints of metabolites compared to genes (Lee, Su, and Huan 2025). Moreover, pathway enrichment results from metabolite measurements in urine or blood plasma can not be directly interpreted as altered pathway biosynthesis, since the enzymes required for many pathways are not present and it is rather to be thought that these metabolites were released into those fluids from organs or blood cells (Lee, Su, and Huan 2025). Here it is also relevant to consider that the origin of the pathways is based on well-established characterization, which has been discussed to hinder novel biological insights (Hu et al. 2025). Hence, we would recommend to always perform literature searches for the top/bottom altered metabolites, especially if they are not well studied. It is also important to take into account that beyond metabolic pathways, metabolites play an active role in gene-regulation by for example binding to receptors and elucidating signalling cascades Farr et al. (2024).

#### Annotated metabolites and coverage in the prior knowledge

There are different points to consider when integrating metabolomics data with PK, which can be divided in several steps:

1. Choose a database for metabolite IDs. Different to gene names, there are no standardized names for metabolites Koistinen et al. (2023), but different databases that collect metabolite information and assign a metabolite ID to each entry, as for example, the Human Metabolome Database (HMDB) with HMDB IDs (Wishart et al. 2007) or the Kyoto Encyclopedia of Genes and Genomes (KEGG) with KEGG IDs (Kanehisa et al. 2017).
2. Investigate metabolite ID mappings. The database often include multiple entries for the same metabolite with different degrees of ambiguity (Pham et al. 2019). Experimentally different degrees of ambiguity are needed due to the machine sensitivity, where for example stereoisomers are not distinguished (Dias et al. 2016). Yet, standard metabolite-sets used as PK often contain one specific metabolite ID, which can hinder mapping to the detected data (Schmidt et al. 2025). If metabolite detection is unspecific due to machine sensitivity, assign all possible metabolite IDs (Table 1) to ultimately increase the overlap with metabolite-sets of interest, translate metabolite IDs, and/or quantify ambiguities, for example by the *MetaProViz* package (Schmidt et al. 2025).
3. Visualize the available metabolite IDs for the detected metabolites using an upset plot and check available metabolite ID types for metabolite classes (Figure 4 A). For instance, for Reactome (Gillespie et al. 2022), which uses ChEBI IDs for pathway enrichment analysis, only a small subset of the detected metabolites will be included, which needs to be taken into consideration when interpreting the results biologically. In order to ensure the maximum overlap between measured metabolites and prior knowledge of choice we recommend assignment of multiple metabolite IDs dependent on the level of ambiguity to the detected metabolites.
4. Given the incomplete coverage in mass spectrometry-based metabolomics and hence the potential low coverage of the PK, adjust the choice of statistical tests. For metabolomics, methods such as ORA should generally perform well since the method offers the ability to restrict the background to the detected metabolites (assay-specific background) (Wieder et al. 2021).

**Figure 4:**
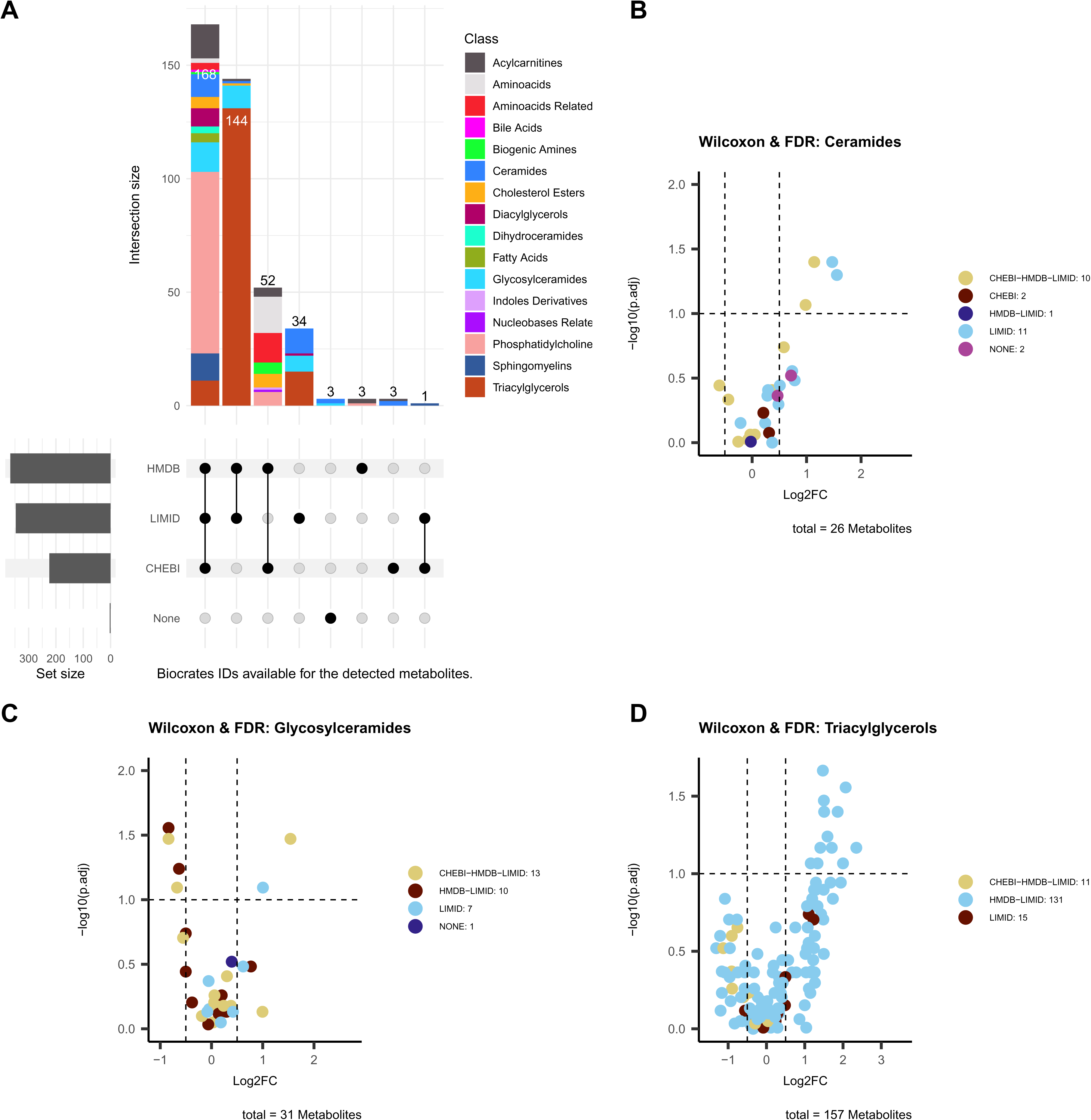
A: Upset plot of the metabolites IDs of the detected metabolites in the Biocrates example dataset colour coded for the Biocrates defined metabolite classes. B-D: Volcano plots of differential analysis comparing TU versus NAT using the Wilcoxon rank-sum test and FDR adjustment, selecting for metabolite panels based on Biocrates class and colour coding for the available metabolite ID types.

**Table 1:** Preview of selected metabolites to showcase ChEBI and HMDB information provided by Biocrates. Biocrates results provide all possible identifiers for each trivial metabolite name. For some metabolites multiple ChEBI and HMDB IDs are available, whilst for other metabolites those databases have no entry.

| Trivial Name | Class | ChEBI | HMDB |
| --- | --- | --- | --- |
| *Ala* | Aminoacids | 15570, 16449, 16977, 76050 | HMDB0000161, HMDB0001310 |
| *C0* | Acylcarnitines | 11060, 16347, 17126 | HMDB0000062 |
| *Cer d16:1/18:0* | Ceramides | None | None |
| *DG 16:0_20:3* | Diacylglycerols | 84402 | HMDB0007111 |

**Table 2:** Compatibility of community detection algorithms with network properties.

| algorithm | undirected | directed | unweighted | weighted |
| --- | --- | --- | --- | --- |
| *cluster_edge_betweenness* | ✓ | ✓ | ✓ | ✓ |
| *cluster_fast_greedy* | ✓ | × | ✓ | ✓ |
| *cluster_fluid_communities* | ✓ | × | ✓ | × |
| *cluster_infomap* | ✓ | ✓ | ✓ | ✓ |
| *cluster_label_prop* | ✓ | ✓ | ✓ | ✓ |
| *cluster_leading_eigen* | ✓ | × | ✓ | ✓ |
| *cluster_leiden* | ✓ | × | ✓ | ✓ |
| *cluster_louvain* | ✓ | × | ✓ | ✓ |
| *cluster_optimal* | ✓ | ✓ | ✓ | ✓ |
| *cluster_spinglass* | ✓ | × | ✓ | ✓ |
| *cluster_walktrap* | ✓ | × | ✓ | ✓ |

#### Available prior knowledge for metabolites

1. Choose a suitable database containing PK:

- For classical mappings with metabolite sets, obtain PK from databases such as *KEGG*, *Reactome* or *WikiPathways*, *RaMP-DB*, *Omnipath*, or *MetSigDB* (*choice of user*):

a. *RaMP-DB* (Braisted et al. 2023) has combined many classical resources trying to account for mismappings between the original resources by using molecular weight. Apart from classical pathway-metabolite sets such as wikipathways, *RaMP-DB* (Braisted et al. 2023) has used *ClassyFire* (Djoumbou Feunang et al. 2016) to assign chemical classes to the metabolites based on their structural components. Chemical class-metabolite sets can be used to understand if certain compound classes are altered in the comparison of interest (Braisted et al. 2023).
b. *Omnipath* Türei et al. (2021) a database of molecular biology PK combines data from more than 1000 resources and incorporates many database clients containing metabolites and utilities for metabolite identifier translation. Apart from pathway-metabolite sets *Omnipath* also offers many other ontology information.
c. To explore signalling function of metabolites by binding to receptors, use *MetalinksDB* (Farr et al. 2024). *MetalinksDB* contains information about metabolite-receptor and metabolite-transporter interaction, as well as whether the binding of the metabolite to a receptor is activating or inhibiting (Farr et al. 2024). Using *MetalinksDB* not only enables combined pathway enrichment analysis on metabolite-gene sets or building a network of those interactions, but also adds biological value by highlighting the signalling potential of the metabolites (Farr et al. 2024).
d. *MetaProViz* includes a collection of annotated metabolite sets usable for classical mapping (Metabolism Signature Database, *MetSigDB*), which includes classical pathway-metabolite sets, chemical class-metabolite sets, and *MetalinksDB* metabolite-receptor and transporter sets, all adapted for classical mapping (Schmidt et al. 2025). Moreover, *MetaProViz* uses the *COSMOS* enzyme-metabolic reaction prior knowledge network (Dugourd et al. 2021) to convert any gene set to gene-metabolite sets, which can be used to perform combined enrichment analysis (Schmidt et al. 2025).
e. When working with human data, the PKN *ocean* includes a manually curated reduced model of human metabolism based on *redHUMAN* of *Recon3D* and has developed a specific adaptation of the weighted mean, using enzyme distance to weight interaction, to perform enrichment analysis (Sciacovelli et al. 2022). *ocean* explores coordinated deregulations of metabolite abundances to generate a metabolic footprint for each metabolic enzyme and ultimately define enzymatic bottlenecks (Sciacovelli et al. 2022).
f. If no metabolomics data are available, but the focus is still on metabolic pathways, use the gene-sets generated by Gaude and Frezza (2016), which include metabolic enzymes and enable weighting for enzymes that are unique for a pathway compared to enzymes present in multiple pathways.
2. (recommended) Exclude certain metabolites from database. The same metabolite can be part of multiple pathways and the database often includes unspecific metabolites like cofactors, ions, H_2_O or CO_2_, which can not be detected with a classical mass spectrometry setup. Many metabolic pathways originate from genome scale metabolic models, which cover the whole reaction in an organism, and those models have not been adapted for the coverage in a metabolomics experiments. We recommend to exclude those metabolites prior to performing enrichment analysis. *MetaProViz* provides pathway-metabolite sets without unspecific metabolites (Schmidt et al. 2025).

#### Biological interpretation of the results

The biological interpretation of the results is crucial and should be done in the context of the biological question and the experimental design. Here it is important to take into account multiple points:

1. Understand the limitations of the PK used, such as potential biases in the underlying original resources. For example, if the PK is based on a specific database, it may not capture all relevant metabolites or pathways. This can lead to biased results and misinterpretation of the data. Moreover, the choice of PK dictates the biological question one can address as discussed in the paragraph above for classical pathway-metabolite sets, chemical class-metabolite sets or receptor-metabolite sets.
2. Understand the coverage of your experimental data in the PK resource. If the PK only includes metabolites that are available in, for example, HMDB, this would in turn exclude many detected ceramides, glycosylceramides and triglycerols, whilst using ChEBI IDs would exclude almost all triacylglycerols (Figure 4 A-D). Here, triacylglycerols include 226 metabolites of which only 11 have a ChEBI ID (Figure 4 D). If the results of the enrichment analysis do not show those pathways as significantly altered, it does not mean that they are not altered, but rather that the PK used is not able to capture them. Consequently, it is important not to over-interpret the importance of the most altered pathways, but rather to use them as a starting point for further investigation not ignoring the detected metabolites that were not covered by the PK resource.
3. (optional) Combine multiple PK resources to increase the coverage of the detected metabolites.
4. Check how many metabolites of e.g. a pathway in the PK have been detected. For example, if a pathway only contains 20 metabolites and 13 of them are detected (65%), this is a stronger evidence than detecting 40 metabolites of a pathway containing 110 metabolites (36%).
5. Understand the impact of metabolite set inflation and deflation on the results. Due to the different metabolite ID types in combination with the metabolite ID coverage in metabolite sets, it sometimes is required to translate metabolite IDs from one identifier type to another. For example, in the Biocrates example data no KEGG IDs are available, yet, if KEGG pathways are used for enrichment analysis, either the KEGG pathways or the metabolite IDs of the data have to be translated, which can lead to one-to-none, one-to-one, one-to-many and many-to-many mappings between the metabolite ID types. This mapping ambiguity can lead to inflation or deflation of the metabolite set size, which needs to be taken into account when interpreting the results. In case of combined enrichment analysis for multiple modalities, take into account the difference in feature space between metabolomics and transcriptomics/proteomics, since this can underestimate the altered metabolites.
6. Take into account the statistical method used for enrichment analysis as it impacts the interpretation of the results (see discussion above).

### Experimental network analysis using statistical methods

Statistical analysis uncovers pairwise relationships between metabolites and/or proteins within each modality. The analysis can be done to assess the cooccurence, association, or regulation of features within a dataset without any PK. The calculation of pairwise coefficients is often done via the *Pearson* or *Spearman correlation* calculation methods, which are useful for exploratory analysis to identify correlated feature pairs, suggesting potential biological relationships or shared pathways (guilt-by-association principle). Other methods may be more suitable depending on the research question, such as *ARACNE* (Algorithm for the Reconstruction of Accurate Cellular Networks), *Bayesian network learning*, *CLR* (Context Likelihood of Relatedness), *GGM* (Gaussian Graphical model), linear regression using *LASSO* (Least Absolute Shrinkage and Selection Operator) regularization, *partial Pearson correlation*, *Random Forest*, or *partial Spearman correlation*. In the resulting network, vertices represent features, such as metabolites or proteins, while edges represent the statistical relationship between them, such as correlation coefficients.

We will showcase the creation of statistical networks based on the implemented functionality of the *MetNet* (Naake and Fernie 2019) package.

1. Choose an algorithm based on the data and biological question:

- *ARACNE* represents an mutual information (MI)-based approach that removes indirect interactions using data processing inequality. The algorithm should be used when aiming to extract the most relevant direct feature interactions from MI-based networks. It is more accurate than pure MI-based methods by reducing false positives by filtering indirect edges. The algorithm requires large sample sizes to estimate MI accurately.
- *Bayesian network learning* infers regulatory networks by modeling conditional dependencies between genes using a probabilistic graphical model. The algorithm may be used when prior biological knowledge is available and probabilistic modeling is needed. The algorithm handles noise well, allows for causal inference, and is able to handle missing data. It may be, however, computationally expensive, requires strong prior assumptions, and may be sensitive to sample size.
- *CLR* uses MI to infer regulatory relationships, normalizing interactions for each feature. The algorithm works well when working with noisy or indirect regulatory influences. It improves MI-based inference by reducing false positives and performs well for global network structures. The algorithm assumes that regulatory interactions follow a Gaussian-like distribution, which may not always be true.
- *GGM* estimates a sparse precision matrix (inverse covariance) to infer direct interactions. The algorithm can be used when assuming a multivariate Gaussian structure in expression data. It identifies direct dependencies and avoids indirect correlations. The algorithm requires large sample sizes to obtain estimates accurately.
- *Linear regression using LASSO regularization* identifies regulatory network by selecting a sparse set of predictor features for each target feature, thereby reducing the number of spurious edges in a network. The algorithm handles high-dimensional data well, reduces overfitting, and provides an interpretable sparse network. The algorithm assumes a linear relationship, may struggle with correlated predictors (features with similar expression profiles, and may be computationally expensive.
- *Pearson correlation* measures linear relationships between feature expression/abundance profiles. The algorithm should be used when linear dependencies exist between feature expression/abundance levels. The algorithm is simple, computationally efficient, and easily interpretable. It does not capture non-linear relationships and is sensitive to noise.
- *Partial Pearson correlation* identifies direct regulatory interactions by controlling for indirect effects. It can be used when confounding influences in a network must be corrected. The algorithm removes indirect correlations, providing clearer direct relationships. It requires large sample sizes to compute reliably.
- *Random forest* infers regulatory network using feature importance scores from an ensemble of decision trees. The algorithm works well when dealing with non-linear relationships and heterogeneous data. It captures complex dependencies, is robust to noise, and does not assume a specific distribution. The algorithm can be computationally expensive and may overfit if not tuned properly.
- *Spearman correlation* captures monotonic (rank-based) relationships between feature expression/abundance levels. The algorithm may be used when the data does not follow a normal distribution or when looking for ranked relationships. It is robust to outliers and captures non-linear relationships. The algorithm is less powerful for detecting direct gene interactions.
- *Partial Spearman correlation* is similar to partial Pearson correlation but for rank-based dependencies. The algorithm controls for confounding variables while dealing with non-linear relationships. It handles non-linearity and indirect interactions. It is less widely used in regulatory network inference and may be computationally demanding.
2. Calculate the adjacency matrix using the *MetNet* package (Naake and Fernie 2019). Specify the algorithms to be run. (optional) Prior to running the model, check if the model tolerates missing values (*NA*) and impute values if needed. (optional) Apply the Benjamini-Hochberg method or any other method for adjustment for p-value adjustment for Pearson and Spearman correlation. The result of each algorithm is stored in a respective assay of the *AdjacencyMatrix* object, an extension of the *SummarizedExperiment* class. Note, that depending on the chosen algorithm, the adjacency matrices are not necessarily symmetric.
3. Examine the distribution of similarity coefficients and p-values, if available (Figure A and B).
4. (optional) Retain highly correlating values (*choice of user*). Feature pairs with low coefficients will be regarded as noise and removed from the adjacency matrix.
5. Create a graph from the consensus adjacency matrix. Specify the graph characteristics, e.g., if the graph should be unweighted *or* weighted or undirected *or* directed.
6. (optional) Remove the singleton components or small networks and obtain the induced subgraph. 5. Visualize network. Results from upstream analyses can be mapped to the network. As an example, the weights of latent factor 1 of the trained MOFA model can be used for colouring of the network vertices (Figure 5 C). Alternatively, we can use logFC of the differential expression analysis. 6. Choose and apply module determination algorithm (*choice of user*). To detect substructures in the graph, module determination algorithms can be applied on the graph and visualize the results (Figure 5 D). Common algorithms are (*igraph* implementation in brackets, see Table 1 for further details):

- *Community structure detection based on edge betweenness* (*cluster_edge_betweenness*) detects communities by iteratively removing edges with the highest betweenness centrality, splitting the network into distinct modules. It works well for small graphs, provides a clear hierarchical structure. The algorithm may be computationally expensive for large networks.
- *Community structure detection via greedy optimization of modularity* (*cluster_fast_greedy*) uses hierarchical agglomeration to merge vertices into communities, optimizing the modularity at each step. The algorithm works efficiently for large graphs, but is less effective for networks with overlapping communities.
- *Community structure detection algorithm based on interacting fluids* (*cluster_fluid_communities*) uses label propagation with a fixed number of communities, allowing them to expand dynamically based on node density. The algorithm is fast. The algorithm requires to predefine the number of communities and may yield inconsistent results across runs.
- *Community structure detection based on Infomap* (*cluster_infomap*) works by simulating random walks on the network and compressing the path description to reveal the best modular structure. The idea is that if a network has well-defined communities, a random walker will spend more time inside a community before jumping to another one. The algorithm captures this by trying to minimize the description length of the walk, effectively grouping vertices that are frequently visited together. The algorithm may be computationally demanding and is sensitive to parameter choices.
- *Community structure detection detection based on propagating labels* (*cluster_label_prop*) assigns labels to vertices, which spread iteratively based on majority voting until convergence. The algorithm scales linearly with the number of vertices and is, thus, works well on large networks. Due to the random nature of initiation, the results can be unstable and inconsistent for certain networks.
- *Community structure detection detection based on leading eigenvector* (*cluster_leading_eigen*) uses spectral clustering by computing the leading eigenvectors of a modularity matrix to find communities. The algorithm may be computationally expensive for large networks.
- *Community structure detection using the Leiden algorithm* (*cluster_leiden*) is a faster version of the Louvain algorithm and yields more accurate solutions. It iteratively refines partitions to ensure well-connected communities by optimizing either the modularity or the Constant Potts model, which does not suffer from the resolution limit. The algorithm may have difficulties finding small communities.
- *Community structure detection by multi-level optimization of modularity* (*cluster_louvain*) optimizes modularity by merging hierarchically small communities into larger ones in a greedy manner. The algorithm scales well to large networks. It may produce different results on each run and may has difficulties finding small communities.
- *community structure detection using the maximum modularity* (*cluster_optimal*) calculates the optimal community structure by maximizing the modularity over all possible partitions. The algorithm guarantees the best modularity score, but is computationally expensive for large network.
- *Community structure detection based on statistical mechanics* (*cluster_spinglass*) models community detection as an energy minimization problem via a spin-glass model and simulated annealing. The algorithm is computationally expensive for large networks and may be sensitive to parameter settings.
- *Community structure detection based on short random walks* (*cluster_walktrap*) simulates random walks on the network to detect community structure, assuming vertices within the same community are more likely to be visited in one walk. The algorithm may be slow for large networks.
7. (optional) Depending on the biological question combine network structure with prior knowledge (Figure 5 E). For example, analyse the module composition via PK resources to check for metabolite classes. Use one-tailed Fisher’s Exact test to check for overrepresentation of a given class.
8. Perform analysis of network structure.

**Figure 5:**
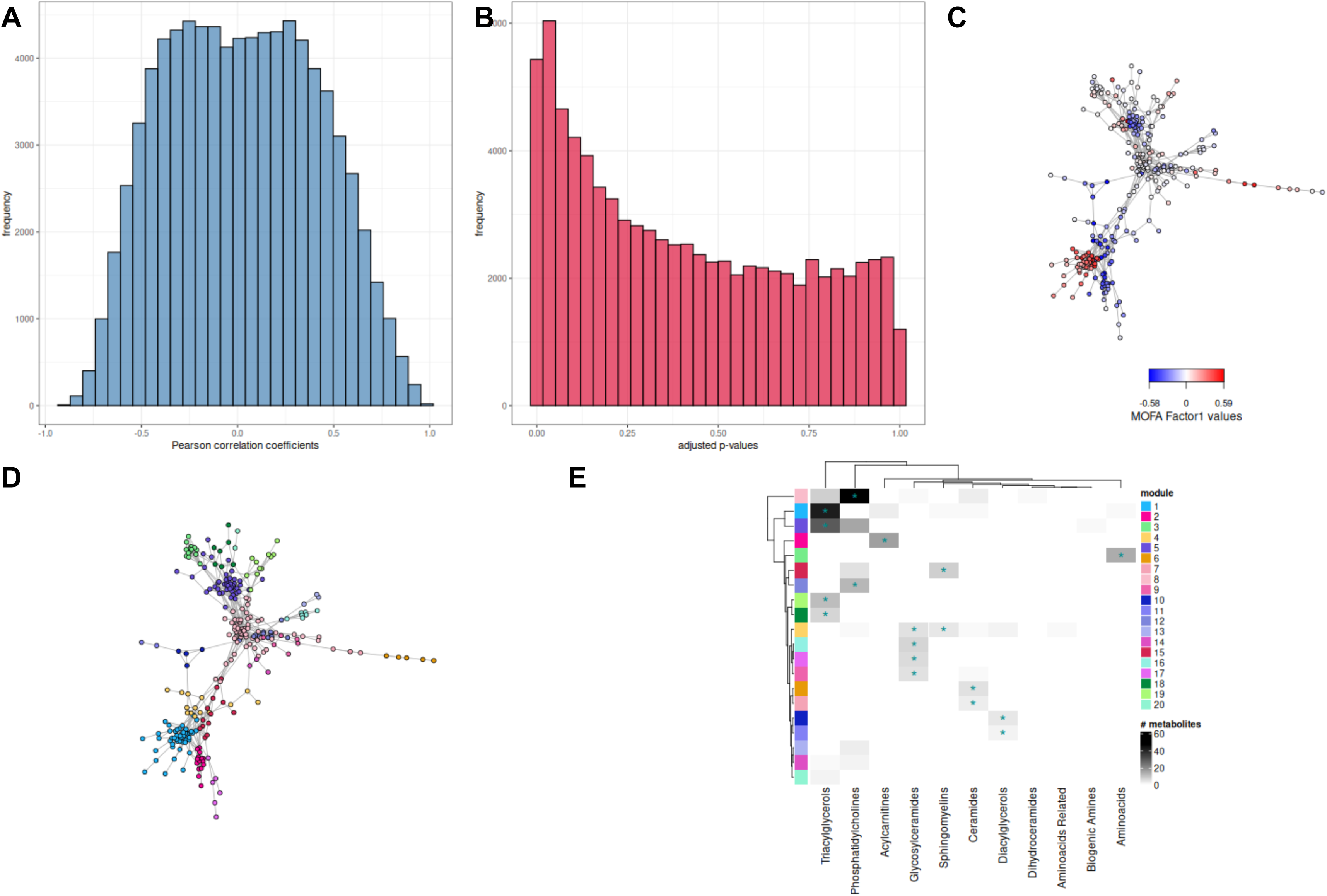
Statistical network analysis of metabolomics data. A: Histogram of coefficients of Pearson correlation-based network. B: Histogram of corresponding adjusted p-values of Pearson correlation-based network using the Benjamini-Hochberg method). C: Thresholded network based on Pearson and Spearman correlation. Each node represents a metabolite and is colored according to its MOFA Factor 1 loading. Edges were retained if the absolute correlation coefficient was > 0.8 (Pearson) and > 0.7 (Spearman), with corresponding adjusted p-values < 0.05 using the Benjamini-Hochberg method. D: Thresholded network based on Pearson and Spearman correlation. Nodes represent metabolites and are colored according by community membership based on the label propagation community detection algorithm. For a legend of module membership refer to panel E. E: Heatmap of chemical class distribution across network modules based on label propagation algorithm. The heatmap shows the abundance of chemical classes (columns) within each module (rows), with color intensity reflecting class count. Asterisks (*) indicate statistically significant overrepresentation of a chemical class within a module (Fisher’s exact test, FDR-adjusted p < 0.05).

## Conclusion

This chapter demonstrated how to perform metabolomics data analysis in *R* using prior knowledge in combination with enrichment analysis or network-based approaches. By integrating tools for preprocessing, differential analysis, enrichment and correlation analysis, and data integration, *R* provides a comprehensive framework for deriving biological insights from metabolomics datasets and combine them with other omics such as proteomics. Parameters such as thresholds, normalization methods, and correction techniques must be carefully chosen to ensure the robustness and reliability of the results.

In order to interpret a metabolomics experiment in its entirety, it is important to understand the limitations of the prior knowledge used, the coverage of the experimental data in the prior knowledge resource, the impact of metabolite set inflation and deflation on the results and the choice of statistical method used for enrichment analysis. Researchers should not solely rely on pathway analysis and often it is more beneficial to perform literature searches for the most altered metabolites, extract metabolite receptors that could be targeted by the altered metabolites or transporters that could release the metabolites into the environment. All of this will require close collaboration between data scientists and experimentalists in the future.

